# Omics Analysis Unveils the Pathway Involved in the Purple Coloration in Tomato Seedling and Fruits

**DOI:** 10.1101/2023.02.09.527816

**Authors:** Rui He, Kaizhe Liu, Shuchang Zhang, Jun Ju, Youzhi Hu, Yamin Li, Xiaojun Liu, Houcheng Liu

## Abstract

The purple tomato variety ‘Indigo Rose’(InR) is favored due to its bright appearance and abundant anthocyanins. *SlHY5* is associated with anthocyanin biosynthesis in ‘Indigo Rose’ plants. However, residual anthocyanins still present in *Slhy5* seedlings and fruit peel indicated there was an anthocyanin induction pathway that is independent of HY5 in plants. The molecular mechanism of color formation in ‘Indigo Rose’ and *Slhy5* mutants is unclear. In this study, we performed omics analysis to clarify the regulatory network underlying coloration in seedling and fruit peel of ‘Indigo Rose’ and *Slhy5* mutant. Results showed that the total amount of anthocyanins in both seedling and fruit of InR were significantly higher than those in *Slhy5* mutant and most genes associated with anthocyanin biosynthesis exhibited higher expression levels in InR, suggesting that *SlHY5* play pivotal roles in flavonoid biosynthesis both in tomato seedlings and fruit. Yeast two-hybrid (Y2H) results revealed that SlBBX24 physically interacts with SlAN2-like and SlAN2, while SlWRKY44 could interact with SlAN11 protein. Unexpectedly, both SlPIF1 and SlPIF3 were found to interact with SlBBX24, SlAN1 and SlJAF13 by yeast two-hybrid assay. Suppression of SlBBX24 by virus-induced gene silencing (VIGS) retarded the purple coloration of the fruit peel, indicating an important role of *SlBBX24* in the regulation of anthocyanin accumulation. These results deepen the understanding of purple color formation in tomato seedling and fruits in an HY5-dependent or independent manner via excavating the genes involved in anthocyanin biosynthesis based on omics analysis.

## Introduction

Anthocyanins comprise a class of primary hydrosoluble pigments belonging to flavonoids, which are widely distributed in plants and confer various coloration in fruit, flower, seed and leaf. Extensive studies have been attracted to plant anthocyanins due to their crucial roles in protecting plants from various biotic and abiotic stressors (i.e., cold, drought, UV irradiation, pathogen), promoting pollination, as well as decreasing the risk of certain types of cardiovascular and neurodegenerative diseases and cancer in human body (Riordan *et al*., 2003; Ma *et al*., 2021; Wang *et al*., 2022).

Anthocyanin biosynthetic pathway have been intensively studied in many species, and various structural genes and transcription factors have been well characterized in a strongly conserved pathway (Liu *et al*., 2018; Outchkourov *et al*., 2018; Butelli *et al*., 2021). The regulation of anthocyanin biosynthesis is controlled by early biosynthetic genes (i.e., phenylalanine ammonia lyase (PAL), 4-coumaryl:CoA ligase (4CL), chalcone synthase (CHS), chalcone isomerase (CHI) and flavanone 3-hydroxylase (F3H)) and late biosynthetic genes (i.e., flavonoid 3’5’-hydroxylase (F3’5’H), dihydroflavonol 4-reductase (DFR), anthocyanidin synthase (ANS), glutathione-S-transferase (GST) and flavonol-3-glucosyltransferase (3GT))(Petroni and Tonelli, 2011). Most genes involved in anthocyanins biosynthesis could be activated or repressed by specific transcription factors as well as controlled by a ternary MBW complex, which consists of basic helix-loop-helix (bHLH), R2R3-MYB transcription factors, and WD40-repeat proteins (Boase *et al*., 2014). In addition, other regulatory factors, such as HY5, ERFs, PIFs and BBXs, WRKYs, also participated in the regulation of anthocyanin biosynthesis (Lloyd *et al*., 2017; An *et al*., 2019; Qiu *et al*., 2019*a*; Xiong *et al*., 2019). Previous studies have proved that *AtBBX21, AtBBX22* and *AtBBX23* induced the accumulation of anthocyanins in Arabidopsis (Datta *et al*., 2007; Chang *et al*., 2008; Xu *et al*., 2016; Zhang *et al*., 2017), while *AtBBX24, AtBBX25* and *AtBBX32* inhibit anthocyanin accumulation (Holtan *et al*., 2011; Gangappa *et al*., 2013; Job *et al*., 2018). *SlBBX20* directly binding the promoter of the anthocyanin biosynthesis gene *SlDFR* to enhanced anthocyanin biosynthesis in tomato fruits (Luo *et al*., 2021). *MdBBX22* induced mdm-miR858 expression via bounding to its promoter, thus governing anthocyanin accumulation in apple (Zhang *et al*., 2022). Additionally, members of B-box (BBX) protein family (i.e., BBX18/20/21/23/24/33) directly conjunct with HY5, cooperatively regulates anthocyanin synthesis in Arabidopsis (Shin *et al*., 2013). Furthermore, *MdWRKY11* could significantly promote anthocyanin accumulation in apple via binding to W-box cis-elements in the promoters of *MdMYB10, MdMYB11*, and *MdUFGT* (Liu *et al*., 2019). Similarly, *SmWRKY44* could activate the expression of *SmCHS, SmF3H* and *SmANS* promoters then positively regulate anthocyanin biosynthesis in eggplant (He *et al*., 2021).

Tomato (*Solanum lycopersicum*) is one of the most consumed vegetable products around the world. In most of cultivated tomatoes, anthocyanins are generally undetectable in fruit. Cultivation attempts have been made to improve anthocyanin content in tomato fruit. ‘Indigo Rose’(InR), a purple tomato variety, which containing the Aft locus and recessive atv locus, exhibits a high-level accumulation of anthocyanin on the fruit peel in a light-dependent manner (Mes *et al*., 2008). Thus, ‘Indigo Rose’ has been frequently taken to underly the molecular mechanism of anthocyanin synthesis in purple tomato fruit (Qiu *et al*., 2016; Sun *et al*., 2019; Wang *et al*., 2020*b*). Previous studies revealed R2R3-MYB transcription factor *SlAN2-like* as an active and critical regulator of anthocyanin biosynthesis, while *SlMYBATV* was identified as the regulatory repressor via competing the binding of *SlAN2-like* to *SlAN1* (Sun *et al*., 2019).

HY5 (Elongated Hypocotyl 5), as a vital regulator of light-dependent development in higher plants, exhibits a dominant function in hypocotyl elongation and lateral root development as well as pigment accumulation (Xiao *et al*., 2022). To date, most of genes and transcription factors involved in anthocyanin regulation were highly associated with HY5. The HY5 protein directly binds to either G-box or ACE-box in the promoters of anthocyanin biosynthetic genes such as *CHS, DFR* then activated their expression, positively regulating anthocyanin accumulation (Shin *et al*., 2007). *CaHY5* can bind to the promoter of *CaF3H, CaF3*′*5’H, CaDFR, CaANS* and *CaGST*, which are well related to anthocyanin biosynthesis or transport, and thereby promote anthocyanin accumulation in pepper hypocotyl (Chen *et al*., 2022). However, residual anthocyanins have been still present in *hy5* mutants of Arabidopsis which indicated that there is an anthocyanin induction pathway which is independent of HY5 in plants (Lee *et al*., 2007; Shin *et al*., 2007). This result has been well proved in *Slhy5* mutants of tomato via analyzing the transcriptome of multiple tissues, which has found eight candidate transcription factors were likely involved in anthocyanin production in tomatoes in an HY5-independent manner (Qiu *et al*., 2019*b*). In the present study, we found that anthocyanins accumulated on the surface of hypocotyls in InR tomato seedling, but not *Slhy5* seedling, at cotyledon emergence. Unexpectedly, residual anthocyanins also accumulated both in the cotyledon and hypocotyls of *Slhy5* seedling which displayed obvious spatiotemporal specificity. Meanwhile, InR fruit peel accumulated large amounts of anthocyanins, particularly in the light-exposing part, while *Slhy5* contained a lower anthocyanin content on the peel of fruit shoulder and no anthocyanins accumulated in the peel of shading part. Therefore, whether other transcription factors substitute or compensate for HY5 to regulate the anthocyanin accumulation?

Advances in transcriptomics and metabolomics play pivotal roles in uncovering complex biological mechanisms of diversified pathways in plants. In this study, we attempted to determine pigment changes in seedling and fruit of tomato (‘Indigo Rose’ (InR) and *Slhy5* mutants) at different developmental stages and excavate the candidate HY5-dependent or independent transcription factors involved in anthocyanin biosynthesis in the seedling and fruit via omics analysis.

## Materials and methods

### Plant materials and growth conditions

The experiment was carried out in an artificial light plant factory in South China Agricultural University. The tomato seeds of wild-type (cv. ‘Indigo Rose’(InR)) and *Slhy5* mutant were generously provided by Dr. Qiu of the College of Horticulture of South China Agriculture University (Qiu *et al*., 2019*a*). Seeds were sanitized in 0.5% sodium hypochlorite for 15 min, then rinsed with distilled water. After being soaked in distilled water for 5 h at 25□, seeds were sowed in sponge cubes (2 cm × 2 cm × 2 cm) with distilled water in the plant growth chamber in the dark at 25□ for 3 days. Then, the seedlings were grown under 300 µmol⋅m^−2^⋅s^−1^ white LEDs (Chenghui Equipment Co., Ltd, Guangzhou, China; 150 cm × 30 cm), 10/14 h light/dark photoperiod, 24 ± 2□, and 65%–75% relative humidity. The seedlings grown for 4th, 5th, 6th and 7th days were sampled and divided into three parts: cotyledon (Except for the samples on the 4th day, which cotyledons still closed), upper 1 cm of hypocotyl, and bottom 1 cm of the hypocotyl (Figure1 a). Three biological replicates were collected for analyses, with each replicate composed of 60 seedlings. Fruits from the *Slhy5* mutants and ‘Indigo Rose’ were sampled at the green-mature stage and fully mature stage with three biological replicates (each replicate consisted of six fruits from different plants). The fruit peel and flesh were carefully split with a scalpel blade. Above-mentioned samples were immediately frozen in liquid nitrogen, and stored at -80□ for further analysis.

**Figure 1.**
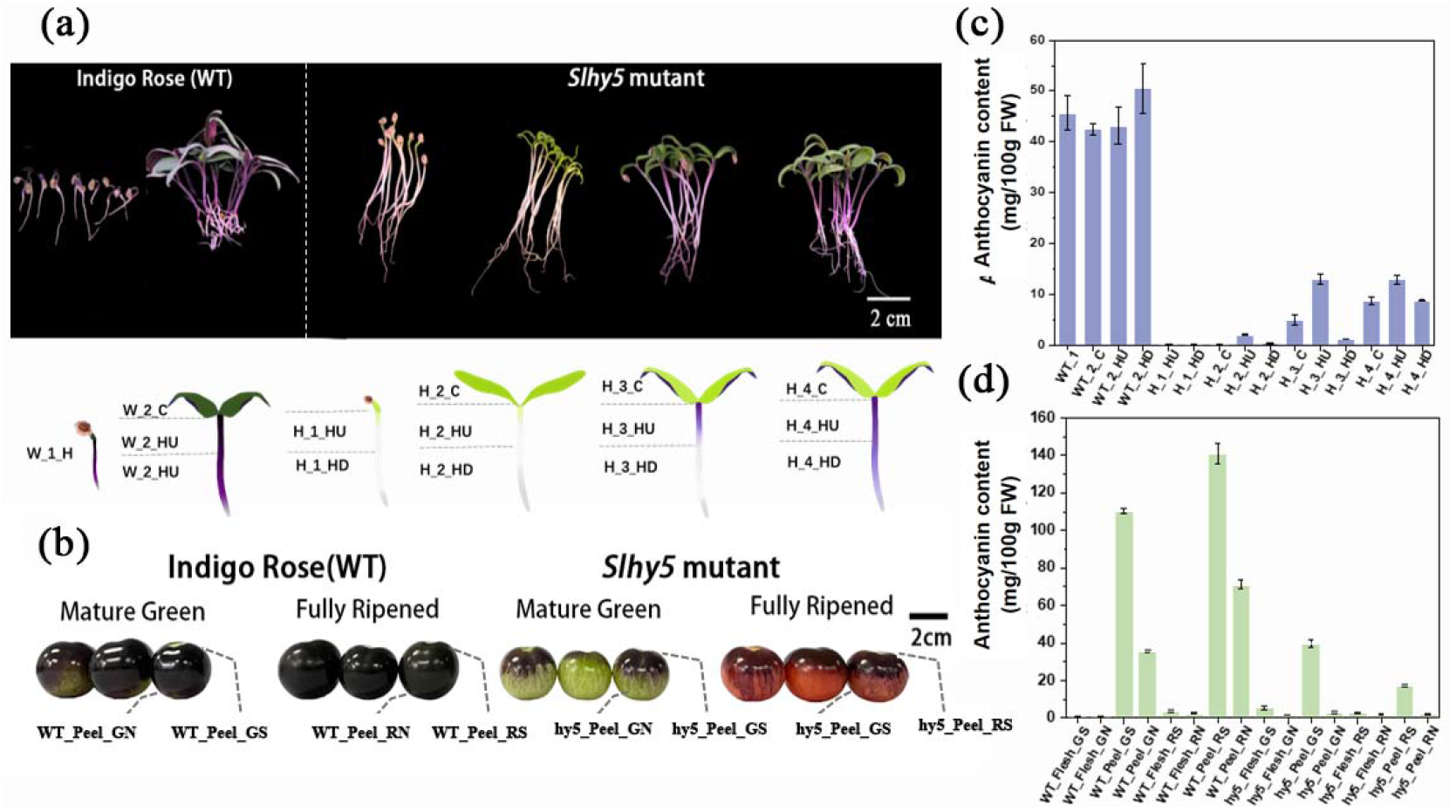
Morphological characterization of InR and *Slhy5* seedling and fruit. (a)Photograph showing the seedling phenotypes of different stages of InR and *Slhy5* seeding for 4-7 days. (b) The phenotype of fruit from InR and *Slhy5* mutants at the mature green-mature stage and fully mature stage. (c) The anthocyanin content of InR and *Slhy5* seedling in different tissues. (d) The anthocyanin content of InR and *Slhy5* fruit s at the green-mature stage and fully ripened stage both in peel and flesh. Error bars indicate SD (n > 3).

### Anthocyanin Assay

The anthocyanin content was performed as described previous study with some modifications (Rapisarda *et al*., 2000). Samples (100 mg) were extracted with 1 mL buffers of pH 1.0 (50 mmol KCl and 150 mmol HCl) and pH 4.5 (400 mmol sodium acetate and 240 mmol HCl) respectively and incubated overnight at 25□. The absorbance of the extract liquor was determined at 510 nm using a UV-spectrophotometer (UV-1600, Shimadzu, Japan).

### RNA Sequencing and Data Analyses

Total RNA was extracted from different parts of the seedling (Figure1 a) and the peel of fruit (Figure1 b) using RNeasy Plant Mini kit (Qiagen, Hilden, Germany). Total amounts and integrity of RNA were assessed using the RNANano 6000Assay Kit of the Bioanalyzer 2100 system (Agilent Technologies, CA, USA). Three independent biological replicates were performed. The RNA-seq sequencing and assembly of seedling and fruit peel were performed by NovoGene Science and Technology Corporation (Beijing, China) and Genedenovo Biotechnology Co., Ltd. (Guangzhou, China), respectively. Total of 3 μg RNA was prepared for sequencing libraries using the NEBNext UltraTM RNA Library Prep Kit for Illumina (NEB, USA) according to manufacturer’s instructions and sequences attributed to each sample by adding index codes. The library preparations were sequenced on an Illumina Hiseq 4000 platform to generate paired-end reads. The raw sequence reads were filtered by removing adaptor sequences and low-quality sequence, and raw sequences were changed into clean reads. Then, the clean reads were then mapped to the tomato reference genome sequence (ITAG 4.0) (https://solgenomics.net/organism/Solanum_lycopersicum/genome/). Padj<=0.05 and |log2(foldchange)| >= 1 were set as the threshold for significantly differential expression. WGCNA analyses were constructed in the BioMarker cloud platform (http://www.biocloud.net).

### Metabolite extraction

Samples of different parts of seedling preparation, extract analysis, metabolite identification and quantification were performed by the novogene database of Novogene Co., Ltd. (Beijing, China). Tissues (100 mg) were individually ground with liquid nitrogen, and the homogenate was re-suspended with pre-chilled 80% methanol, and 0.1% formic acid by vortexing. The supernatant was injected into an LC-MS/MS system. UHPLC-MS/MS analyses were performed using a Vanquish UHPLC system (ThermoFisher, Dreieich, Germany) coupled with an Orbitrap Q ExactiveTM HF mass spectrometer (ThermoFisher). Structural analysis of metabolites was determined using standard metabolic operating procedures. MRM was used to conduct metabolite quantification. All metabolites identified were subject to partial least squares discriminant analysis. Principle component analysis (PCA) and Orthogonal Partial Least Squares Discriminate Analysis (OPLS-DA) were carried out to identify potential biomarker variables. For potential biomarker selection, variable importance in projection (VIP) ≥ 1 and fold change (FC) ≥ 2 or ≤ 0.5 were set for metabolites with significant differences.

### Yeast Two-Hybrid Assay

The amplified full-length CDSs of *SlPIF1, SlPIF3, SlAN2-like, SlAN2, SlAN11, SlAN1, SlHY5, SlJAF13, SlBBX24, SlWRKY44* were amplified and inserted into pGADT7 and pGBKT7 vectors, respectively. Primers used for amplified and plasmid construction are listed in Supplementary Table S1. Different combinations of bait and prey vectors were cotransformed into Y2Hgold then cultured on SD/-Leu-Trp (SD-LT) medium supplemented at 28 □ for 2–3 days. 2.5 μL of aliquots were then patched on SD/Ade/His/Leu/Trp plates with 5-bromo-4-chloro-3-indolyl-α-D-galactopyranoside (X-α-Gal) and incubated at 28 □ for 3 days.

### Virus-induced SlBBX24 gene silencing in tomato

Specific coding regions SlBBX24 fragment were selected for VIGS vector construction by the VIGS design tool (http://solgenomics.net/tools/vigs). A 300 bp fragment of the coding region of SlBBX24 was amplified using the forward primer (5′-gtgagtaaggttaccgaattc ATGAAGATACAGTGTGATGTG-3′) and the reverse primer (5′-cgtgagctcggtaccggatcc AGTGGCTAAGAAGCGTTGGTG-3′). The PCR product was ligated the sequence into pTRV2 vector to construct the TRV2::SlBBX24 vector. The recombinant plasmid and TRV1 vector were transferred into Agrobacterium GV3101. Resuspensions of pTRV1 and pTRV2 (as a negative control) or its derivative vectors were mixed at a 1:1 ratio and then infiltrated into a mature green stage tomato. The injected fruits were grown in a growth chamber at 22 ± 2 □ and 12 h of light/12 h of darkness, and relative humidity was controlled in the range of 70 ± 5%. After two weeks, fruits were harvested and stored at −80□ for RNA and qRT-PCR analysis to assess the degree of silencing.

### RNA extraction and Quantitative Reverse Transcription PCR

Total RNA was extracted from different parts of the seedling and the peel of fruit using RNAex Pro Reagent (Accurate Biotechnology Co., Ltd., Hunan, China), and its quality and quantity were evaluated using a Nanodrop ND-1000 spectrophotometer (Thermo Fisher Scientific). The first cDNA strand was synthesized using Evo M-MLV RT for PCR Kit (Accurate Biotechnology Co., Ltd., Hunan, China). The relative expression levels were determined by performing a quantitative reverse transcription PCR (qRT-PCR) analysis using a LightCycler 480 system (Roche, Basel, Sweitzer) with an Evo M-MLV RT-PCR kit (Accurate Biotechnology Co., Ltd., Hunan, China). The PCR thermocycling protocol was as follows: 95□ for 30 s, followed by 40 cycles of 95□ for 15 s and annealing for 60□ for 30 s. Three biological replicates and three qRT-PCR technical replicates were performed for each sample. *SlUBI* was used as the reference gene for normalization. The primer sequences used for the qRT-PCR analyses are listed in Table S1.

## Results

### Morphological characterization of InR and Slhy5 seedling and fruit

On the stage which hypocotyl emergence and cotyledons still close, the surface of hypocotyls of InR seedlings exhibited purple color whereas *Slhy5* seedlings display white color (hardly accumulate anthocyanins) and longer hypocotyl (Figure 1a and Figure S1). Subsequently, a considerable anthocyanins accumulation was observed in cotyledons and hypocotyls of InR seedlings (Figure 1a and Figure S1). Interestingly, *Slhy5* seedling developed an opposite phenotype that seedling displayed lower anthocyanin and exhibit spatiotemporal character of anthocyanin production, which anthocyanin accumulated first in the upper part of the hypocotyls then gradually developed in the lower part in a light-dependent manner (Figure 1a and Figure S1). The anthocyanin content both in cotyledons and hypocotyls of *Slhy5* seedling was lower than those of InR seedling, respectively (Figure 1c and Figure S1).

Anthocyanin content just accumulated in peels at the green-mature stage and fully mature stage in InR and *Slhy5* fruits, and few anthocyanins were found in the fruit flesh (Figure 1b,d and Figure S2). Additionally, higher anthocyanins accumulated in light-exposing peel part of InR fruits than shading part (Figure 1b,d). Unexpectedly, residual anthocyanins have been still present in the peel of *Slhy5* fruit shoulder, though the anthocyanins contents were lower than InR fruit peel. However, anthocyanins accumulation was hardly detected in the shading peel part of *Slhy5* fruit (Figure 1b,d). These observations suggested that *Slhy5* had a predominant role in tomato pigmentation in a light-dependent manner and there might be some regulators controlling anthocyanin biosynthesis in an HY5-independent manner.

### Changes of metabolites and genes in the cotyledon of InR seedlings and Slhy5 seedlings

The color of cotyledon and hypocotyl were transformed continuously from green to purple during seedling development in InR seedlings and *Slhy5* seedlings. To explore the metabolites and genes changes during InR and *Slhy5* seedlings development, metabolic and transcriptome analysis of cotyledon, the upper and lower part of the hypocotyl were carried out at different developmental stages respectively (Figure 1a). PCA was performed on detected metabolites to demonstrate the similarity in metabolic profiles among the samples (Figure S3). In the two-dimensional PCA plot, three biological replicates of each sample tended to group, indicating the high reproducibility of the generated metabolome data (Figure S3). Many metabolites and genes in the seedling varied considerably in terms of different tissues of different variety. Therefore, variation of metabolites and genes in three parts (cotyledon, upper and lower part of the hypocotyl) of InR seedlings and *Slhy5* seedlings was investigated, respectively.

Four metabolites groups were observed in cotyledon based on the level of annotation of metabolites similarity. Group 1, which metabolites levels of InR seedlings cotyledon (W_2_C) were obviously higher than those in *Slhy5* seedlings (H_2_C, H_3_C and H_4_C), including tulipanin, rutin, butin etc., were belong to flavonoids, flavones and flavonols, anthocyanins according to KEGG analyses (Figure S4). So, anthocyanins and flavonoids might be responsible for the distinction of purple coloration of cotyledon between InR and *Slhy5* seedlings.

The transcriptome data validated the authenticity and accuracy of the metabolic analysis. A weighted gene co-expression network analysis (WGCNA) was performed on the genes of cotyledon. A slight relationship with purple coloration was displayed in the green module (Figure 2b), and the genes related to this module were annotated according to KEGG pathway enrichment analysis, which involved the flavonoid biosynthesis pathways (Figure 2c). Anthocyanins metabolism-related transcriptional factors (i.e., *SlAN1, SlAN2*, SlAN2-like) and the structural genes (i.e., *SlPAL, Sl4CL, SlCHS, SlCHI, SlDFR, SlANS*) displayed higher expression levels in the cotyledon of InR seedlings than *Slhy5* seedlings (Figure 2c,d). Meanwhile, the expression of these genes increased with the development of *Slhy5* seedlings cotyledon, which is consistent with the increasing trend of flavonoid and anthocyanins contents (Figure 2c,d).

**FIGURE 2.**
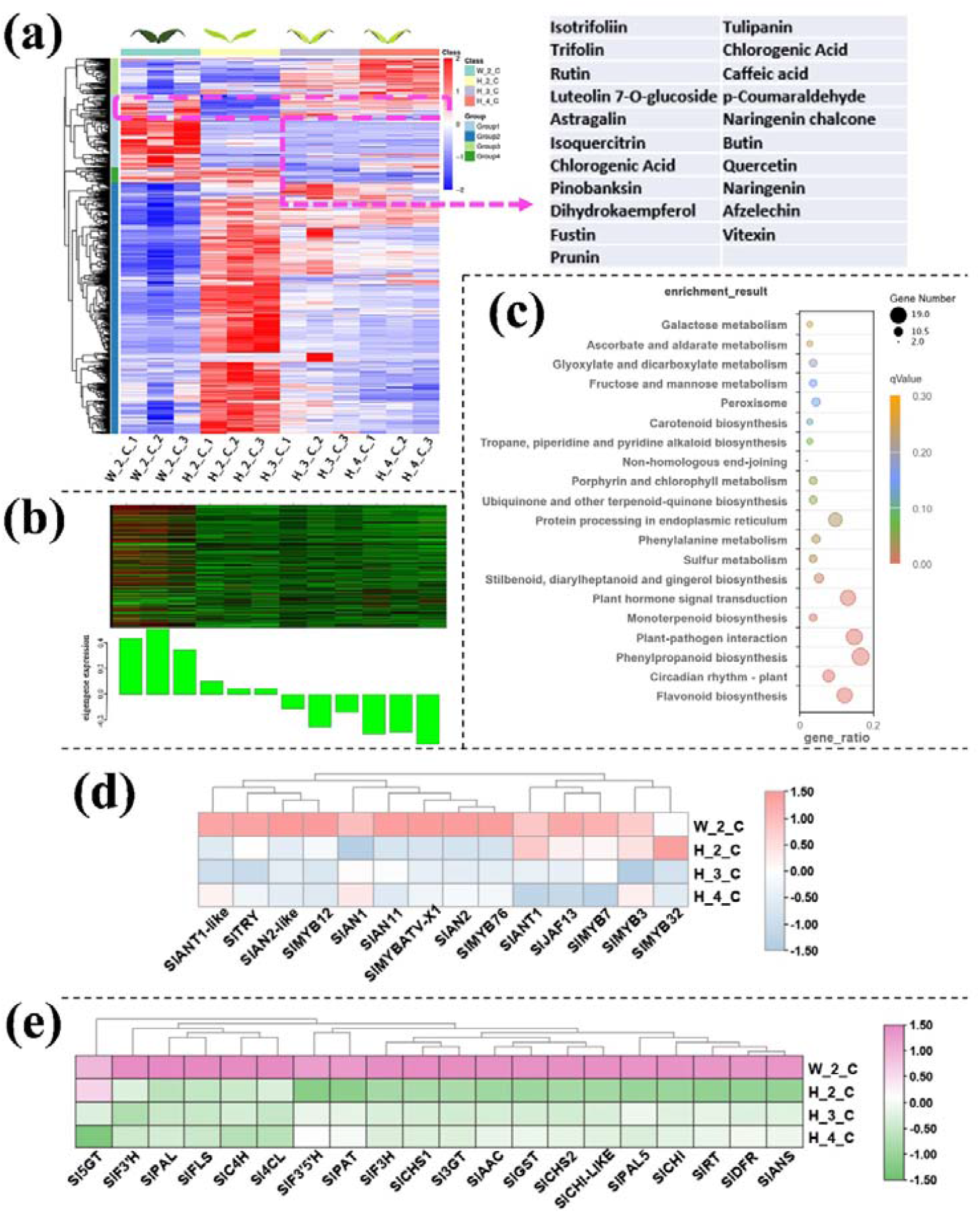
Differentially accumulated metabolites (DAMs) accumulation pattern consisted with the color variations during seedling development of cotyledon (a). The genes in green module that consisted with the color variations during seedling development of cotyledon (b). Top 20 enriched KEGG pathway enrichment of the genes in green module (c). The FKPM values of the transcriptional factors (d) and the structural genes (e) related to flavonoid and anthocyanin biosynthesis.

### Changes of metabolites and genes in the upper part of the hypocotyl of InR seedlings and Slhy5 seedlings

In the upper part of hypocotyl, the differential metabolites in group 1 which positively correlated and have similar consistent patterns with the purple coloration in the hypocotyl, also contained a variety of flavonoids such as petunidin 3-O-glucoside, tulipanin, isotrifoliin (Figure 3a). The KEGG enrichment analysis displayed that the terms ‘Flavone and flavonol biosynthesis’, ‘Photosynthesis’, ‘Anthocyanin biosynthesis, ‘Vitamin B6 metabolism’, ‘Starch and sucrose metabolism’ were significantly enriched in the Group 1 (Figure S5). WGCNA identified genes in the black module with significant co-expression with the metabolites biosynthesis in the flavonoid pathway, which were responsible for the purple coloration observed in the upper part of hypocotyl (Figure 3b). KEGG enrichment analysis were performed on the genes in the black module and showed that these DEGs were enriched mainly in flavonoid biosynthesis pathways (Figure 3c). Consistent with the data from the transcriptomic analysis, both the anthocyanin positive regulatory genes, such as *SlAN2, SlAN2-like, SlAN1, SlAN11* (except for the negative regulatory genes *SlMYB7, SlMYB3, SlMYB32*), and the anthocyanin biosynthetic genes, including *SlPAL, Sl4CL, SlCHS, SlCHI, SlF3H, SlF3*′*H, SlDFR, SlANS* and *Sl3GT* (Figure. 3d,e), exhibited much higher expression in the upper part of InR seedlings hypocotyl than those of *Slhy5* seedlings. Compare with H_1_HU and H_2_HU, the genes involved in anthocyanins under H_4_HU and H_3_HU displayed higher expression levels (Figure. 3 d,e).

**FIGURE 3.**
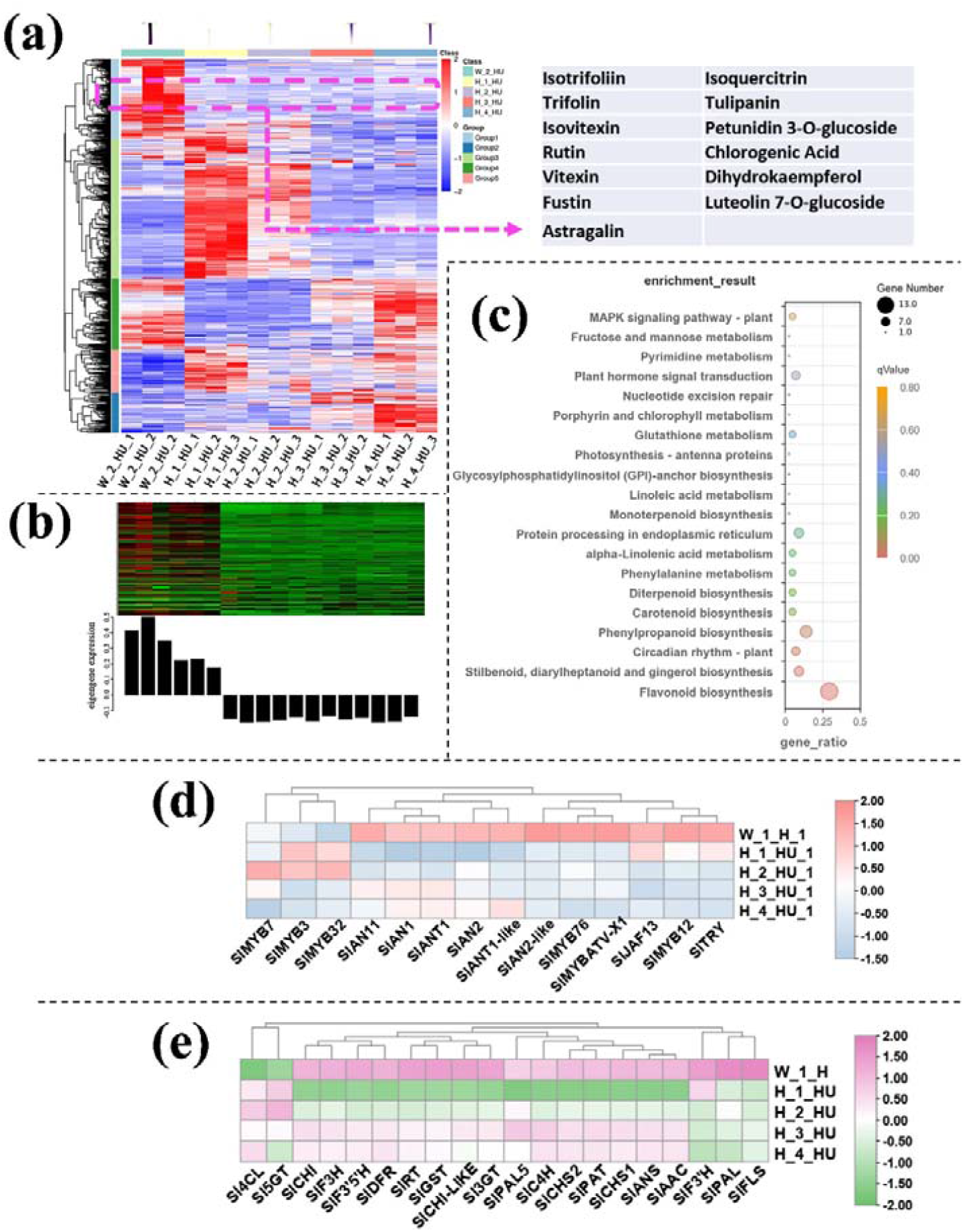
Differentially accumulated metabolites (DAMs) accumulation pattern consisted with the color variations during seedling development of the upper part of hypocotyl (a). The genes in green module that consisted with the color variations during seedling development of upper part of hypocotyl (b). Top 20 enriched KEGG pathway enrichment of the genes in black module (c). The FKPM values of the transcriptional factors (d) and the structural genes (e) related to flavonoid and anthocyanin biosynthesis.

### Changes of metabolites and genes in the lower part of the hypocotyl of InR seedlings and Slhy5 seedlings

Similarly, annotated metabolites in the lower part of the hypocotyl could be divided into several large groups based on similar variation tendency, respectively. Among these metabolites, the group 1 that metabolites present in higher levels in InR seedlings than *Slhy5* seedlings, which were consisted with the color variations during seedling development. These metabolites including petunidin 3-O-glucoside, tulipanin, isotrifoliin (Figure 4a). KEGG analysis showed that these metabolites in different parts of the seedling were mainly enriched in flavone and flavonol biosynthesis, and anthocyanin biosynthesis (Figure S6). The genes in paleturquoise module according to WGCNA analysis of transcriptomic data from the lower part of hypocotyl were consistent with the increasing trend of anthocyanin contents (Figure 4b). The KEGG enrichment analysis showed that the terms ‘flavonoid biosynthesis’ pathway was significantly enriched (Figure 4c). Meanwhile, the expression levels of anthocyanin biosynthesis genes in the lower part of InR hypocotyl were higher than *Slhy5* (Figure 4d,e). In addition, the expression of most anthocyanin biosynthetic genes in the lower part of the *Slhy5* seedling hypocotyl under different stages exhibited the following trend: H_4_HD ≈ H_3_HD > H_2_HD > H_1_HD.

**FIGURE 4.**
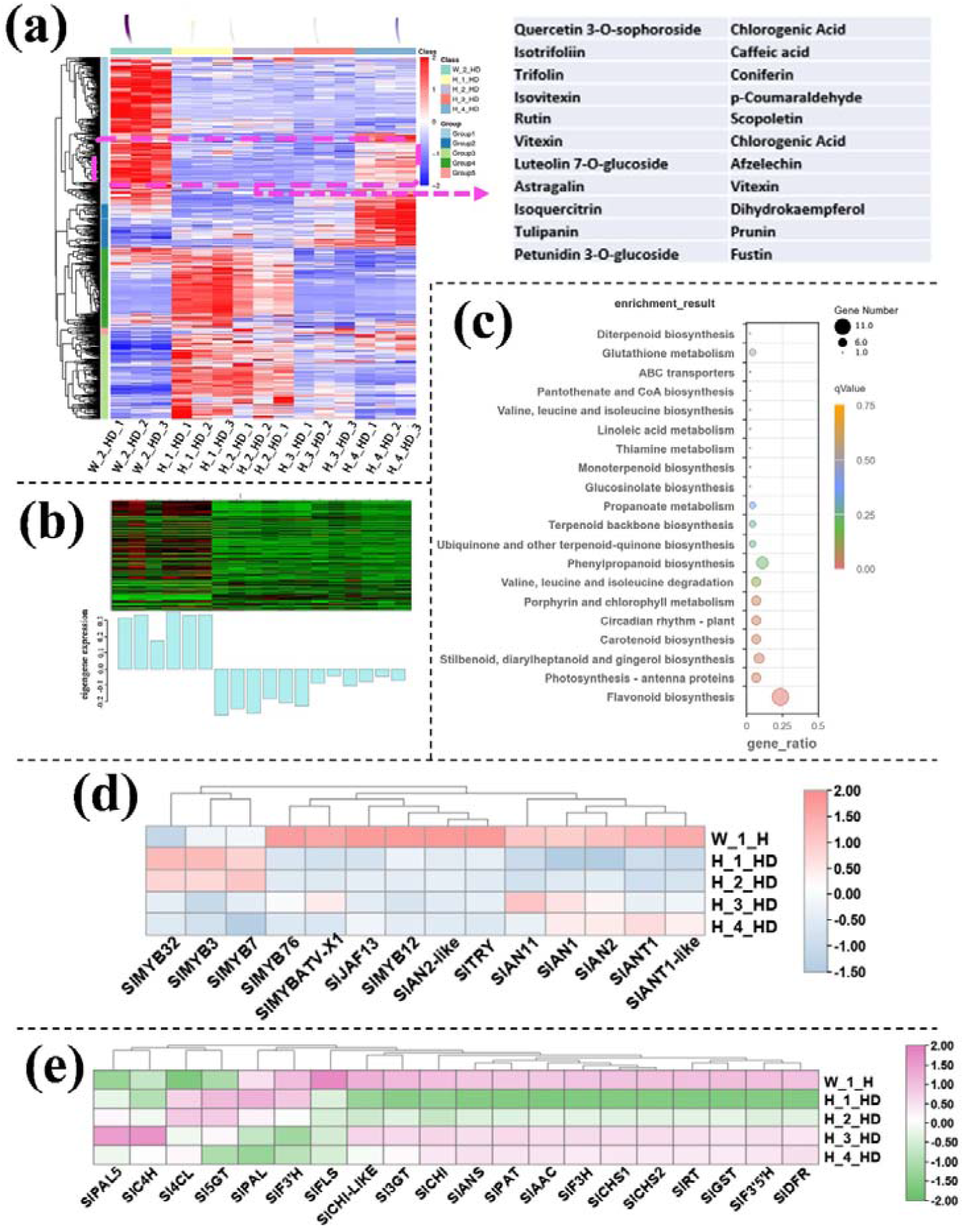
Differentially accumulated metabolites (DAMs) accumulation pattern consisted with the color variations during seedling development of the lower part of hypocotyl (a). The genes in green module that consisted with the color variations during seedling development of lower part of hypocotyl (b). Top 20 enriched KEGG pathway enrichment of the genes in Paleturquoise module (c). The FKPM values of the transcriptional factors (d) and the structural genes (e) related to flavonoid and anthocyanin biosynthesis.

#### 3.5 Changes of metabolites and genes in different parts of *Slhy5* seedlings

Based on WGCNA analysis, we identified a cyan module whose gene expression pattern was associated with the phenotype of anthocyanin synthesis in *Slhy5* seedling at the third and four development stages (Figure 5a, b). 21 co-expressed genes in the cyan module were significantly correlated with pigment accumulation in *Slhy5* seedlings. These indicated that bHLH (*SlAN1*) was a hub gene involved in the positive regulation of flavonoid metabolism in *Slhy5* seedlings (Figure 5c), possibly by affecting structural node genes, such as *SlCHI, SlCHS, SlAN3, SlDFR* and *SlRT* etc.

**FIGURE 5.**
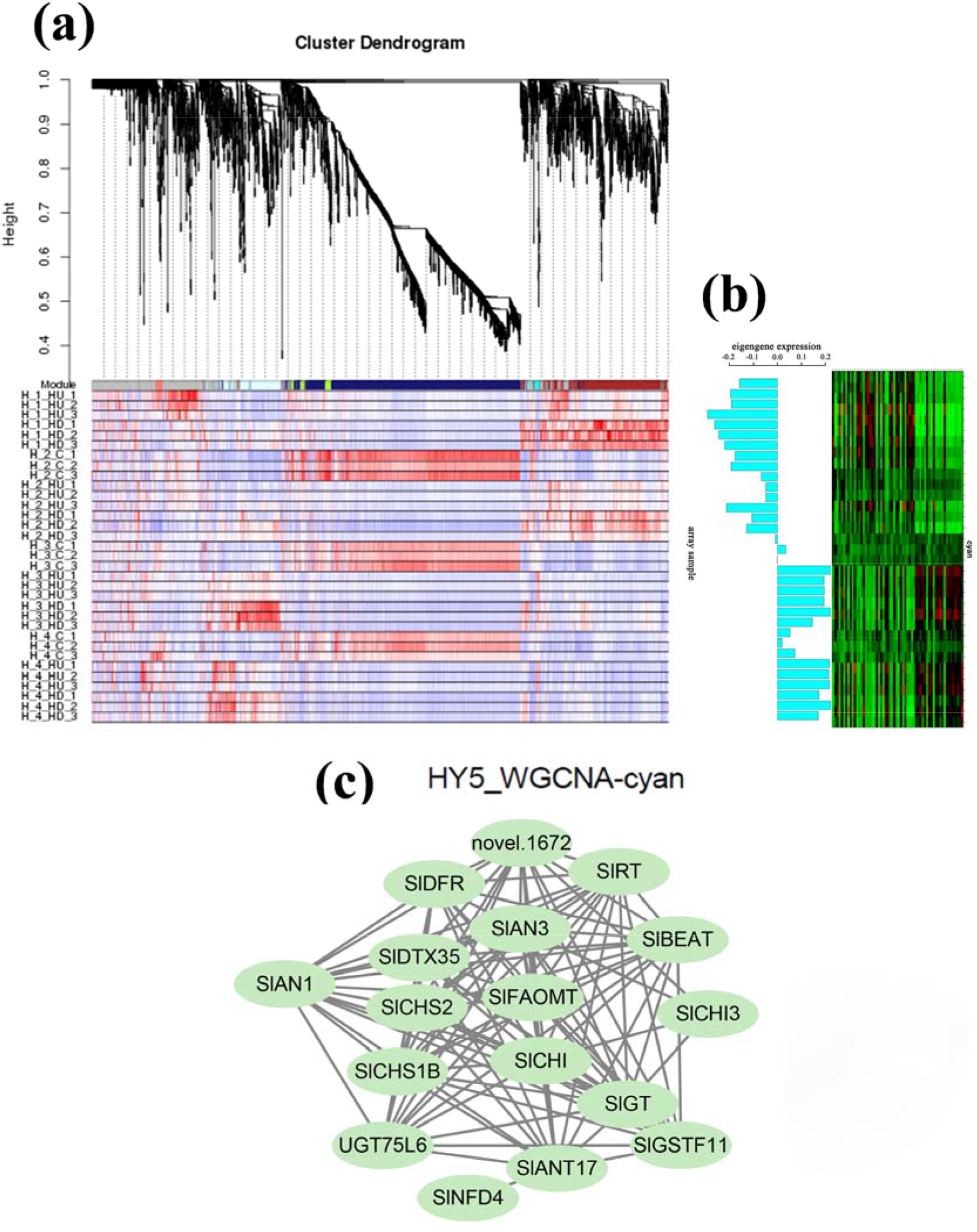
Hierarchical cluster tree showing seven modules obtained by WGCNA in *Slhy5* seedlings (a). Differential expression of genes in the accumulation pattern consisted with the color variations during seedling development of Slhy5 mutant seedlings (b). Interaction network of DEG in the cyan model in Slhy5 seedling (c).

### Screening of differentially expressed genes of tomato fruit

Similar to seedlings, residual anthocyanin production was also observed in the peel of *Slhy5* fruit shoulder, whereas anthocyanin was almost undetectable in the shading part of *Slhy5* fruit (Figure1 b,d). To underly gene expression changes over the fruit peel of InR and *Slhy5*, RNA-Seq analysis was conducted. The number of differentially expressed genes had a very high variance among InR and *Slhy5* fruit peel. Regarding WT-S-vs-slhy5-S (purple-colored peel of InR fruit compared to purple-colored peel of *Slhy5* fruit), a total of 2995 DEGs including 1360 up- and 1635 down-regulated genes were detected (Figure S7). These genes were enriched in KEGG pathways related to photosynthesis, carbon fixation in photosynthetic organisms, phenylpropanoid biosynthesis, phenylalanine metabolism, flavonoid biosynthesis and flavone and flavonol biosynthesis (Figure 6a), which the gene FKPM values in InR fruit peel was higher than those of *Slhy5* (Figure 6c,d). These results well demonstrated the importance of the HY5 in the color formation (both green and purple) of tomato fruits.

**FIGURE 6.**
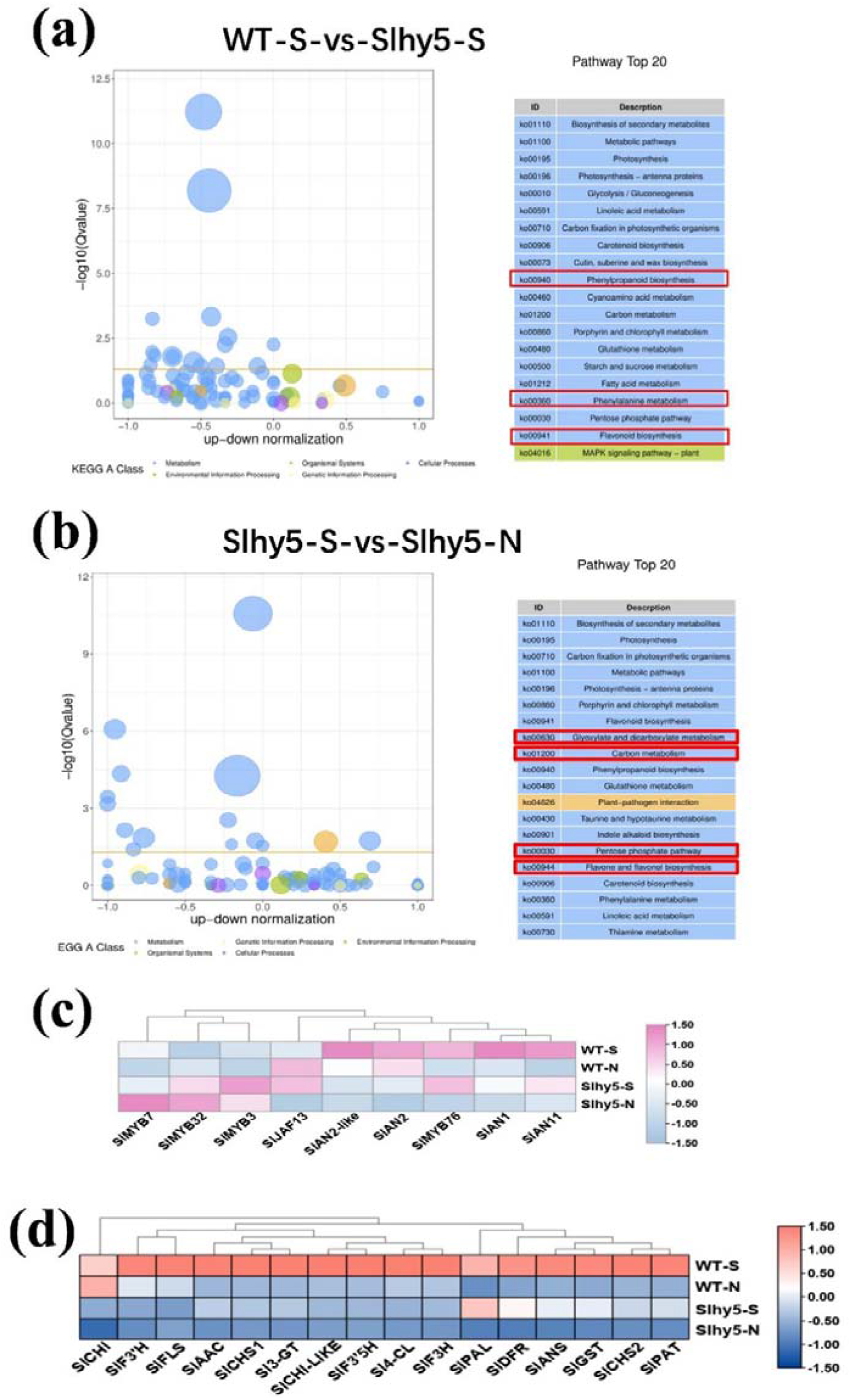
KEGG enrichment of differential genes in WT-S vs. Slhy5-S (a) and Slhy5-S vs Slhy5-N (b). The FKPM values of the transcriptional factors (c) and the structural genes (d) related to flavonoid and anthocyanin biosynthesis.

Meanwhile, to further investigate the DEGs related to *Slhy5* fruit peel coloration, we compared the FPKM values of Slhy5-S-vs-Slhy5-N (purple-colored peel compared to white-colored peel in *Slhy5* fruit). A total of 2393 DEGs were identified from the groups of Slhy5-S-vs-Slhy5-N (Figure S7). According to KEGG enrichment analysis, the 20 top-ranked pathways contributed by these DEGs were photosynthesis, carbon fixation in photosynthetic organisms, porphyrin and chlorophyll metabolism, flavonoid biosynthesis, phenylpropanoid biosynthesis, flavone and flavonol biosynthesis, carotenoid biosynthesis and phenylalanine metabolism. A set of genes involved in flavonoid metabolism in the light-exposing part of the *Slhy5* fruit peel displayed a higher expression level than the shading part (Figure 6b). Therefore, we predicted that the flavonoid biosynthesis pathway was leading to the purple peel coloration in *Slhy5* fruit peel, that was proved that other genes or TFs were involved in anthocyanin biosynthesis in a HY5 independent manner.

To further verify expression patterns of the DEGs in seedlings and fruit, the genes *SlAN1, SlPAL, SlCHS, SlCHI, SlF3H, SlDFR, SlANS, Sl3-GT, SlAAC, SlGST* for qRT-PCR verification (Figure S8). The qRT-PCR results were consistent with the transcriptomic analysis results.

### The genes involved in the MYB-bHLH-WD40 (MBW) complex that activates anthocyanidins in tomato fruit

To explore the regulation of flavonoids metabolism, the possible interaction of flavonoids metabolism-related genes was tested utilizing yeast two-hybrid (Y2H) assays. SlPIF1, SlPIF3, SlAN2-like, SlAN2, SlAN11, SlAN1, SlHY5, SlJAF13, SlBBX24, SlWRKY44 were selected for the candidate genes based on the RNA Sequencing results which might be contribute to anthocyanin biosynthesis. We found that SlBBX24 physically interacts with SlAN2-Like and SlAN2, but not with SlAN11 in yeast, while SlWRKY44 could interact with SlAN11 protein only, and showed no affinity for SlBBX24. Additionally, these results indicated that SlBBX24 could interact with SlPIF1 and SlPIF3 but not SlPIF4. Unexpectedly, SlAN1 and SlJAF13 displayed the same physical interaction which could interacted with both SlPIF1 and SlPIF3.

### SlBBX24 may be involved in anthocyanin accumulation in tomato fruit peels

To investigate whether the biosynthesis of anthocyanin is regulated by candidate genes, a target gene of *SlBBX24*, was silence by VIGS approach. Compared to the empty vector control, the expression of *SlBBX24* was reduced at 7 days after infiltration, and anthocyanin was less vibrant than that of control fruits. (Fig.7). qRT-PCR analyses were performed to test the expression changes of the genes involved in anthocyanin biosynthesis. As shown in Figure 7, silencing of *SlBBX24* inhibited the expression levels of anthocyanin structural genes, including EBGs (*SlC4□ and SlCHS1*) and LBGs (*SlDFR* and *SlANS*) as well as the positive regulators (*SlAN1, SlAN2, SlAN2-like, SlAN11, SlHY5*, and increased the expression levels of negative regulators (*SlMYBATV, SlTRY* and *SlMYB76*). Therefore, it suggests that *SlBBX24* might be a positive regulator of anthocyanin biosynthesis in tomato fruit.

**FIGURE 7.**
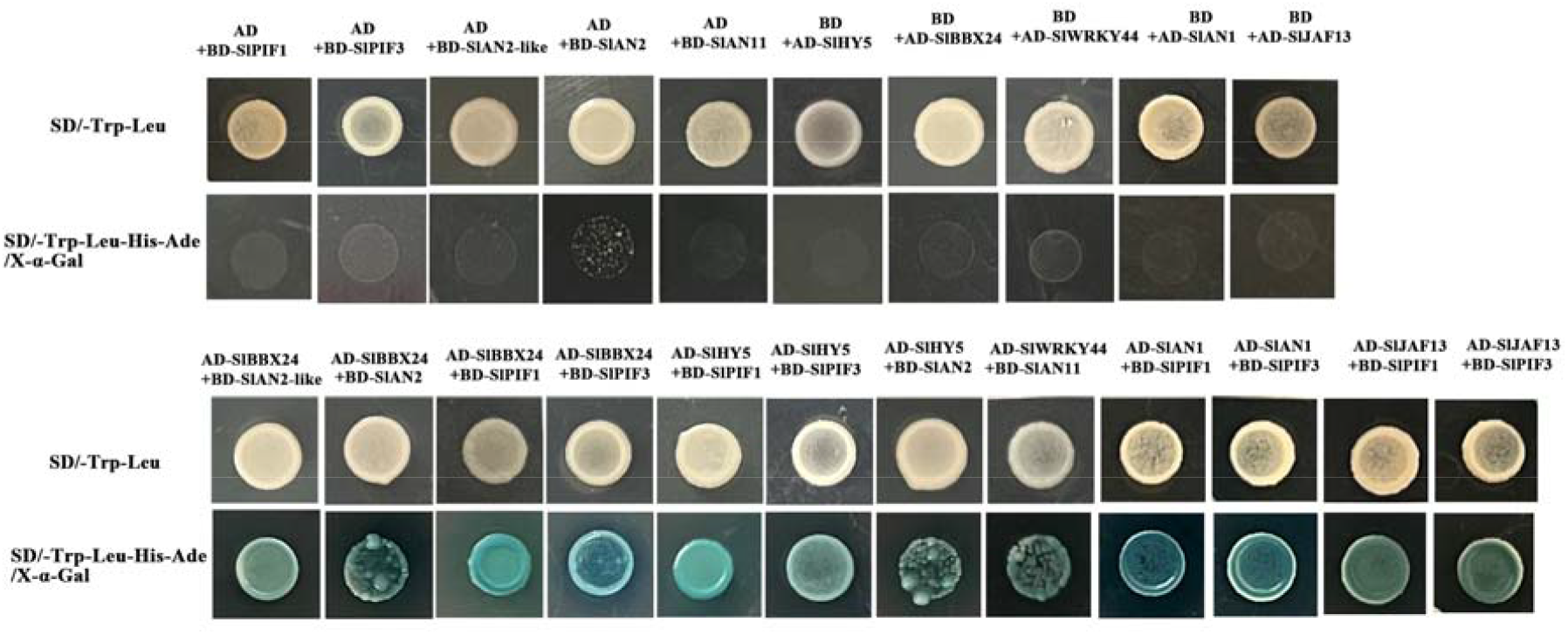
Yeast two-hybrid assay of the protein interactions between candidate genes involved in the components of the MBW complex that activates anthocyanidins in tomato fruit.

## Discussion

### Flavonoids might be attributed to major color differences among InR and Slhy5 mutant seedlings and fruit peel

In recent years, integrated metabolomic and transcriptomic analyses as efficient tools to underly the molecular mechanisms of key metabolic pathways in plants (Rothenberg *et al*., 2019; Wang *et al*., 2020*a*; Jiang *et al*., 2022; Light and Stress, 2022). A detailed description of secondary metabolic changes occurring in the whole germinated seeds as well as cotyledons, hypocotyls and roots from 3 to 9 days old tomato seedlings via LCMS profiling, which provided a new perspective to study metabolic networks controlling flavonoid biosynthesis in tomato (Roldan *et al*., 2014). The metabolite variants across 20 major tomato growth tissues and stages were explored via combining transcriptome and metabolome approaches, which verified novel transcription factors that controlling steroidal glycoalkaloids and flavonoids pathways (Li *et al*., 2020). In the present study, an integrated analysis of the transcriptome and metabolome was conducted to reveled the differences between wild-type (InR) and *Slhy5* seedlings at different growth and develop stages. Total 987 metabolic components were accumulated specifically in InR and *Slhy5* seedling, of which amino acids and organic acids, flavanones, flavones, isoflavonoids accounted for a large proportion (Table S2). Flavonoids, a product of the phenylpropanoid metabolism pathway, are extensively distributed in numerous plants which are composed of various subclasses, including flavanones, flavones, isoflavonoids, anthocyanins, and flavonols. Anthocyanins as a key flavonoid subgroup, responsible for pigmentation in flower, fruit, seed, and leaf (Liu *et al*., 2018). Our metabolic profiling found metabolites in different parts of the seedling were mainly enriched in flavone and flavonol biosynthesis (Figure 2). Meanwhile, levels of flavonoids including anthocyanins in InR seedling were obviously higher than those of *Slhy5* (Table S2). A KEGG analysis in WT-S-vs-Slhy5-S, Slhy5-S-vs-Slhy5-N were related to phenylpropanoid biosynthesis, phenylalanine metabolism, flavonoid biosynthesis and flavone and flavonol biosynthesis (Figure 5a). These genes were enriched in KEGG pathways related to photosynthesis, carbon fixation in photosynthetic organisms, phenylpropanoid biosynthesis, phenylalanine metabolism, flavonoid biosynthesis and flavone and flavonol biosynthesis (Figure 5a), and the gene FKPM values in InR fruit peel was higher than those of *Slhy5*. These results were well indicated that main pigment components in InR and *Slhy5* seedlings and fruit peel were flavonoids.

### SlHY5 acts as a master regulator to control anthocyanin biosynthetic in seedlings and fruit of tomato

Anthocyanin biosynthetic genes are regulated directly by the MBW complex consisting of MYB, bHLH and WDR proteins. Aft and Atv are two important loci that well associated with anthocyanin biosynthesis in tomato. MYB TFs occupy the major determinant position in the control network of anthocyanin biosynthesis and have been well demonstrated (Chen *et al*., 2019; Yan *et al*., 2021). Four R2R3 MYB TF genes (*SlAN2, SlANT1, SlANT1-like* and *SlAN2-like*), were previously identified to regulate anthocyanin biosynthesis in tomato (Schreiber *et al*., 2012; Kiferle *et al*., 2015; Meng *et al*., 2015). Two bHLH TFs, *SlAN1* and *SlJAF1*3, were recently reported to regulate anthocyanin production in tomato (Qiu *et al*., 2016; Cao *et al*., 2017). Otherwise, a tomato WDR protein, *SlAN11* was also involved in anthocyanin synthesis (Gao *et al*., 2018).

HY5 well characterized as a positive regulator of anthocyanin synthesis in a light-dependent manner. Knock-down *SlHY5* transcription significantly reduced the anthocyanin levels both in seedlings and fruit of tomato (Qiu *et al*., 2019*a*). *HY5* activate the expression of *PAP1* expression via directly binding to G-and ACE-boxes in the promoter region, positively induced the accumulation of anthocyanin in Arabidopsis (Shin *et al*., 2013). In apple, *MdHY5* involved in the regulation of anthocyanin biosynthesis via positively regulated both its own transcription and that of *MdMYB10* by binding to E-box and G-box motifs, respectively (An *et al*., 2017). Consistently, *PyHY5* alone or interacted with *PyBBX18* activate the expression of PyMYB10 and PyWD40 which subsequently regulate the anthocyanins accumulation in red pear (Wang *et al*., 2020*c*). In this study, expression levels of *SlAN2-like, SlAN2, SlAN1* and *SlAN11* were higher in InR than *Slhy5* both in seedling and fruit, and the expression pattern of these genes were consistent with pigment accumulation (Figure 2,3,4,6). Two-hybrid (Y2H) assays determined *SlHY5* regulated anthocyanin biosynthesis through interacted with *SlAN2* (Figure 7), consistent with the result that *SlHY5* as a positive regulator of anthocyanin biosynthesis in vegetative tissues of tomato (Zhi *et al*., 2020). Therefore, *SlHY5* might be the master regulator to control anthocyanin accumulation in InR seedling and fruit via mediating the transcriptional activity of an MBW complex and the enhanced expression of key genes, such as *SlCHI, SlCHS, SlF3H, SlDFR* and *SlANS*.

### Possible regulatory mechanisms of anthocyanin biosynthesis in an HY5-independent manner in tomato

Besides MYB and bHLH, other TF families also regulate anthocyanin biosynthesis. *SlBBX20* could directly bind to the promoter of *SlDFR* to activate its expression, thus promoting anthocyanin accumulation in tomato (Luo *et al*., 2021). In apple, *MdBBX37* interacted with *MdMYB1* and *MdMYB9*, two key positive regulators of anthocyanin biosynthesis, and inhibited the binding of those two proteins to their target genes and, therefore, negatively regulated anthocyanin biosynthesis (An *et al*., 2019). *PpBBX18* and *PpBBX21*, antagonistically regulate anthocyanin biosynthesis via competitive association with *PpHY5* in the peel of pear fruit. Also, discoveries have emphasized the importance of WRKY protein in the control of flavonoid pathway and its relationship to the MBW complex (Lloyd *et al*., 2017). *MdWRKY11* was able to bind to W-box cis-elements in the promoters of *MdMYB10, MdMYB11*, and *MdUFGT* then regulated anthocyanin synthesis in apple flesh (Liu *et al*., 2019). In present study, different from InR seedling, some TFs were detected in *Slhy5* seedlings by RNA-Seq, such as WRKYs, BBXs, and NACs might compensate for the function of HY5 and contributed the expression of related genes involved in anthocyanin synthesis. Y2H assays revealed that SlBBX24 could interact with SlAN2-like, and SlAN2, which likely have positive function in the regulation of anthocyanin biosynthesis (Figure 7). Also, SlWRKY44 also could interact with SlAN11 protein (Figure 7), which is consistent with the result that the WRKY factor physically interacts with the AN11 in yeast two-hybrid analysis (Quattrocchio *et al*., 2006). Silencing of *SlBBX24* via virus-induced gene silencing (VIGS) led to the downregulation of the expression of structural genes and caused a decrease in anthocyanin accumulation (Figure 8). Additionally, PIFs play a role in the biosynthesis of plant pigments. *PIF3* could specifically bind to the G-box element of anthocyanin biosynthesis□related structural genes promoter to up-regulate anthocyanin accumulation in a HY5-dependent manner under far-red light (Shin *et al*., 2007). Furthermore, *PIF4* negatively regulated anthocyanin accumulation by inhibiting *PAP1* transcription in Arabidopsis seedlings (Qin *et al*., 2022). In this study, SlPIF1 and SlPIF3 could physically interacts with SlAN1 and SlJAF13, as well as SlBBX24. We speculated that *SlPIF1, SlPIF3, SlBBX24 SlWRKY44* might involve in anthocyanin biosynthesis in a manner independent or dependent of *SlHY5*. Taken together, given the PIFs and MBW were the key regulators of anthocyanin biosynthesis, we propose a model to clearly illustrate this mechanism (Figure 9). In InR, *SlHY5* expression was induced by light and then activating the activity of MBW complex to regulate the anthocyanin accumulation. While in *Slhy5* seedling and fruit, PIFs or several other transcription factors might involve in coordinating anthocyanin biosynthesis, like BBXs, WRKYs. More thorough and rigorous molecular studies should be performed to explore the relationship of PIFs or other transcription factors which might involve in anthocyanin biosynthesis.

**FIGURE 8.**
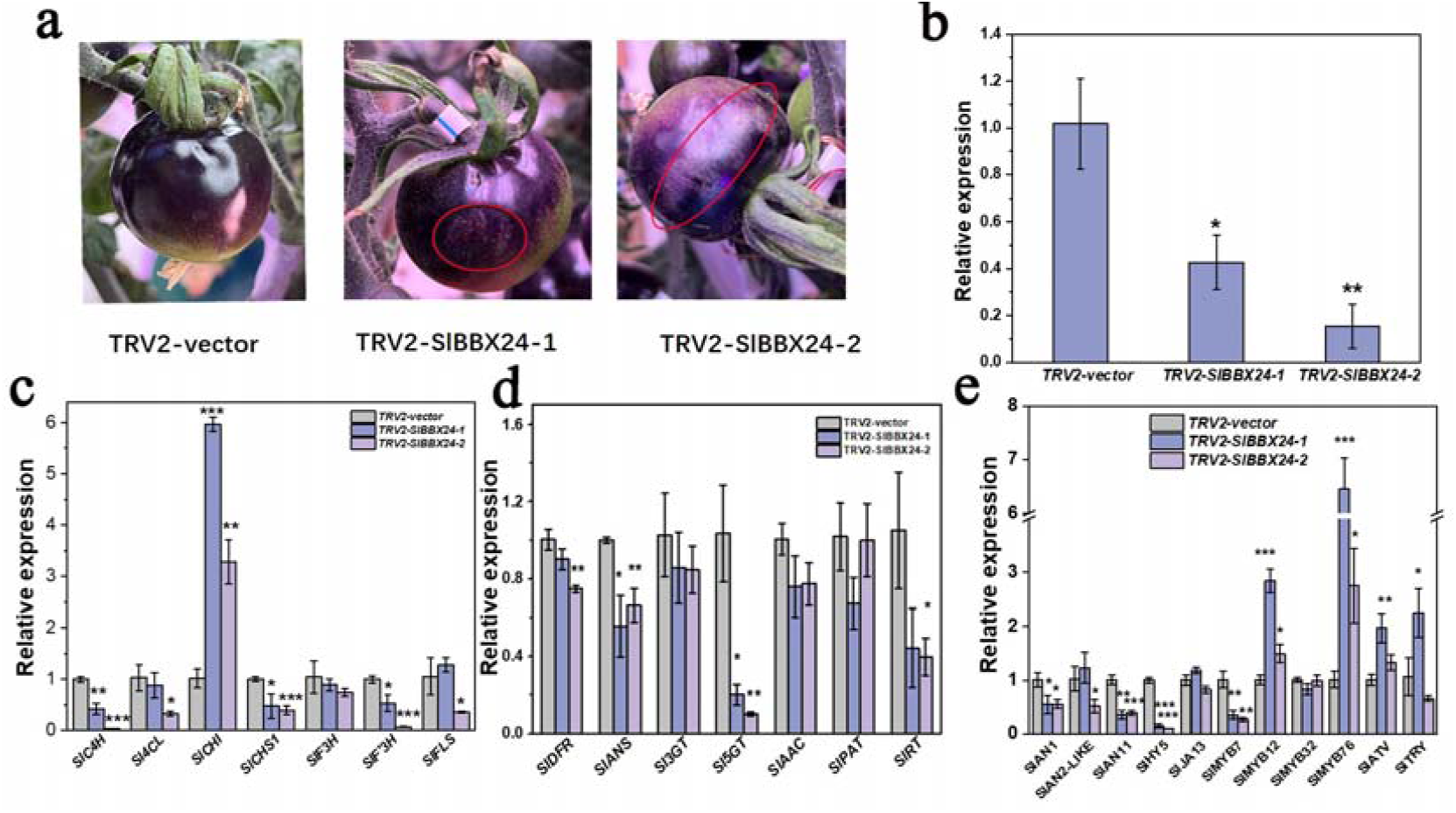
Analysis of VIGS of the *SlBBX24* gene. (A) Suppression of *SlBBX24* by virus-induced gene silencing (VIGS) retarded the purple coloration of the tomato fruit peel. (B) Relative expression of *SlBBX24* in fruit peels of TRV2-inoculated and TRV2-SlBBBX24-inoculated for 7 days. (C-E) Relative expression of anthocyanin structural genes of fruit peels of TRV2-inoculated and TRV2-*SlBBX24*-inoculated for 7 days. Statistically significant differences between purple zone and green zone determined by Student’s t-test. (** p < 0.01, * p < 0.05).

**Figure 9.**
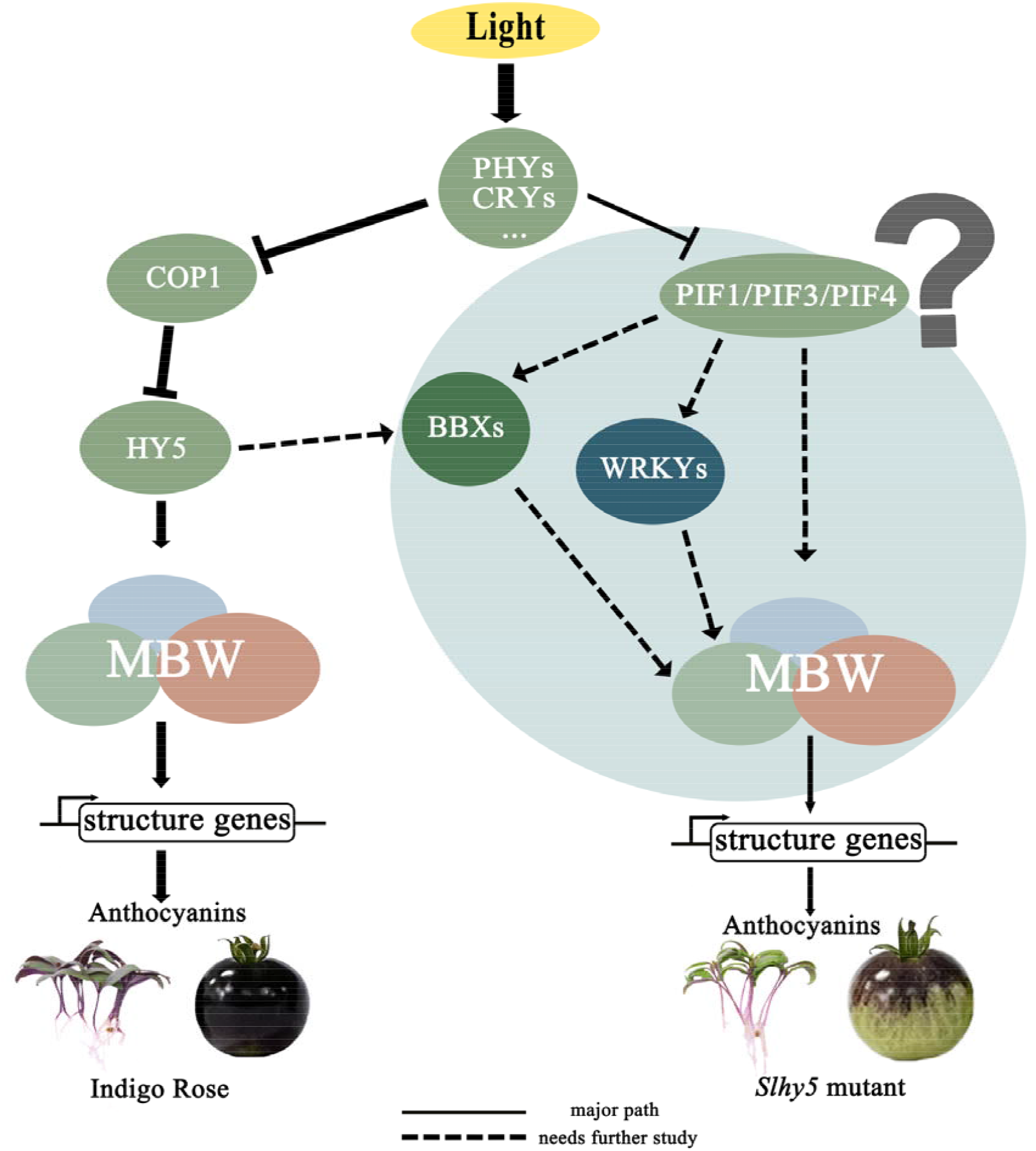
A putative model showing the anthocyanin induction pathways that might be dependent or independent of HY5 in tomato.

## Conclusion

HY5 has a pivotal role in regulating anthocyanin accumulation in tomato. *Slhy5* mutant were created via the CRISPR/Cas9 system from ‘Indigo Rose’. *Slhy5* mutants displayed significantly lower anthocyanin accumulation than InR both in seedling and fruit. Interestingly, detectable levels of anthocyanins are present in hy5 mutant seedling and fruit, and the pigment accumulation displayed obvious spatiotemporal specificity in the *Slhy5* seedling. These results indicated that other regulators exist to regulate anthocyanin biosynthesis in an HY5-independent manner. The total amount of flavonoids in InR was significantly higher than in *Slhy5* mutant and most of the genes associated with anthocyanin biosynthesis displayed higher expression levels in InR. SlBBX24 likely regulate anthocyanin biosynthesis by interacting with SlAN2-like, SlAN2, while SlWRKY44 interacting with SlAN11. Moreover, SlPIF1 and SlPIF3 seemed to involve in anthocyanin biosynthesis through interacting with SlBBX24. We identified *SlBBX24* as a target to silence to produce less anthocyanins in tomato fruit peel, indicating an important role of *SlBBX24* in the regulation of anthocyanin accumulation. These results deepen the understanding of purple color formation in tomato seedling and fruits in an HY5-dependent or independent manner via excavating the genes involved in anthocyanin biosynthesis based on omics analysis.

## Supplementary data

Additional Supporting Information may be found in the online version of this article.

**Figure S1**. Cross sections of cotyledons and hypocotyls of InR and *Slhy5* seedlings.

**Figure S2**. Phenotype of the fruit from the tomato cultivar ‘Indigo Rose’(InR) and Slhy5 mutants at the fully mature stage.

**Figure S3**. Principal component analysis result of all samples analyzed for metabolite contents.

**Figure S4**. Bubble diagram of the enriched KEGG pathways among the differentially abundant metabolites of Group 1 in the cotyledon of InR seedlings and *Slhy5* seedlings.

**Figure S5**. Bubble diagram of the enriched KEGG pathways among the differentially abundant metabolites of Group 1 in the upper part of InR seedlings and *Slhy5* seedlings.

**Figure S6**. Bubble diagram of the enriched KEGG pathways among the differentially abundant metabolites of Group 1 in the lower part of InR seedlings and *Slhy5* seedlings.

**Figure S7** Bar graph of differentially expressed genes (DEGs) in InR and .*Slhy5* fruit peel.

**Figure S8**. Expression pattern validation of genes involved in anthocyanin biosynthetic in tomato seedling and fruit peel using RT-qPCR verified the RNA-Seq results.

**Figure S9**. KEGG pathway enrichment analysis performed in H_1_HD.vs.W_1_H.

**Table S1**. Primers used in this study.

**Table S2**. Metabolites in ‘Indigo Rose’ and the Slhy5 seedling at different stages.

## Author contributions

RH and HL designed the project. RH, KL, JJ, YH, and SZ performed experiments. RH, HL, YL, and XJL analyzed the data. RH, KL and HL wrote the article.

## Data availability

All data referred to are included in the main text or Supporting Information of this manuscript.

